## Supplementary information for "Non-diffusive slow heat dissipation induces high local temperature in living cells"

### Supplementary Notes

#### Supplementary Note 1 | Fast fluorescence lifetime ($\tau_{FAST}$) of FPT for determination of temperature change in living cells

In the TCSPC used in this method, the fluorescence emission time after excitation, which is obtained by measuring the arrival time of photons of FPT detected synchronously with the excitation pulse laser (20 MHz), is integrated by repeated scanning (Figure 1b). In the conventional analysis of fluorescence lifetime, the average fluorescence lifetime ($\tau_f$) is obtained from the value of each component obtained by approximating the fluorescence decay curve drawn using a large number of photons integrated over a long time (60 s) with a two-component exponential function. In contrast, the fast lifetime ($\tau_{FAST}$) is simply determined from averaging the average arrival time of the obtained photons (Figure 1b). It should be noted that the parameters obtained from conventional fluorescence lifetime determination (e.g., the number of components, their respective values, and their proportions) contain photochemical information about the fluorophore, which is useful for detailed investigation of the chemical environment surrounding the fluorophore.

The $\tau_f$ of FPT is slightly longer than the $\tau_{FAST}$ because elimination of the excitation pulse-derived signal (instrumental response function, IRF) was performed by limiting the approximation range (fluorescence decay curves below 2.5 ns were excluded) when calculating the conventional fluorescence lifetime.

We investigated the acceptable variability in fast lifetime determination under limited photon availability when measuring small intracellular temperature changes (with a temperature resolution of approximately 0.2 °C)[1] using FPT. First, the photon count obtained in the most photon-limited high-resolution temperature measurement conducted in this study was 16,967 ± 4,209 ($n$ = 30 cells). In contrast, the temperature response curve for fast lifetime measurements (Figure 1c) indicates that approximately 10,000 photons are required to determine $\tau_{FAST}$ with variance below the temperature resolution of about 0.2 °C (corresponding to a change of about 0.1 ns in the S.D. value of the fast lifetime; Figure 1f). Thus, the variability in FPT fast lifetime determination does not affect the measurements in this study. Second, the number of photons originating from dark noise in the detector is 8.7 ± 1.7 ($n$ = 10 fields of view), which gives an error to $\tau_{FAST}$ of approximately 0.01 ns. This indicates that the error introduced by dark noise is negligible.

### Supplementary Note 2 | Confirmation of temporal and spatial resolution of intracellular temperature measurement by the fast fluorescence lifetime ($\tau_{FAST}$) of FPT

In high-speed temperature tracking, the response time of FPT can affect its spatio-temporal resolution. Because the time required for the conformational change of FPT has been unknown, we investigated this by using a controllable artificial heat source. Polyacrylamide polymer, the main backbone of FPT, exhibits a temperature-dependent phase transition above the lower critical solution temperature (LCST), causing FPT to respond within a specific temperature range (Supplementary Figure 8a). During the process of temperature relaxation to below the LCST of FPT after the heating stops, the fluorescence lifetime change of FPT is determined almost entirely by its structural relaxation, not by the temperature relaxation (Supplementary Figure 8a). In the temperature tracking experiments of this study, the base temperature and the temperature during heating were set within the range where the FPT is temperature-responsive (Supplementary Figure 8a). Here, cells cultured at a low temperature (13 °C) at which FPT is not temperature-responsive, were heated by an IR laser to a temperature (31 °C) at which FPT is temperature-responsive. Supplementary Figure 8b shows the rapid relaxation of the fast lifetime of FPT in cells measured with high temporal resolution (every 9 ms) after the heating was stopped. Given that the first frame after the fluorescence lifetime drop does not reflect the temperature, the result indicates that the time of structural relaxation of FPT obtained both in solution and in cells are equal to or less than the time of scanning (9 ms). This value was used to discuss the temporal (**a**) and the spatial (**b**) resolution of temperature measurement using FPT.

#### (a) Temporal resolution

The parameters related to the temporal resolution include the fluorescence lifetime of FPT (~10 ns); the scanning time for fluorescence lifetime imaging (9 ms); the time required for photon integration, which is calculated as 1.2 ms based on the number of photons (approximately 10,000 photons) required to determine the fluorescence lifetime with an accuracy of 0.1 ns (Figure 1f) and a photon acquisition efficiency of $8.2\times10^3$ ms$^{-1}$; and the response time of FPT (< 9 ms). Among these factors, the scanning time for fluorescence lifetime imaging (9 ms) is the rate-limiting step, which determines the temporal resolution of this intracellular temperature measurement.

#### (b) Spatial resolution

The error (e.g., spatial resolution) with respect to spatial information in temperature mapping using FPT is determined by the diffraction limit of light in microscopy or the distance traveled within the response time of diffusing FPT (as described above, less than 9 ms: Supplementary Figure 8). The molecular diffusion time of FPT in cells was quantified by the fluorescence recovery after photobleaching (FRAP) method ($D_{FPT}$ = 0.13 μm$^2$ s$^{-1}$: Supplementary Figure 9). The distance that FPT moves by diffusion within the time required for the structural transition of FPT upon temperature change was calculated from the following equation:

$$\Delta x^2 = 4Dt \qquad \text{(S1)}$$

where $x$ is the displacement distance, $D$ is the diffusion constant, and $t$ is the time. Because this calculated distance (below 0.07 μm) is smaller than the diffraction limit of light (0.28 μm), the spatial resolution is determined by the diffraction limit of light.

### Supplementary Note 3 | Analysis of time constants for intracellular temperature relaxation

The time course of the intracellular temperature 70 msec after stopping continuous heating in Figure 3c was approximated by the following equation:

$$\Delta T(t) = \sum_{i=1}^{N} \left[A_i exp\left(-\frac{t}{\tau_i}\right) + C_i\right] \qquad \text{(S2)}$$

where $A_i$, $\tau_i$ and $C_i$ represent amplitude, time constant and $\Delta T$ after infinite time of components $i$ respectively. Calculations with increasing values of $N$ until the $t$ test value > 2 was no longer satisfied detected a maximum of two components of relaxations: the measured 231 ± 1.53 msec (47.6%) and 3.33 ± 0.141 sec (17.9%), and relaxation earlier than the tracking limit accounted for 34.5% ($\Delta T$ after infinite time was 0).

### Supplementary Note 4 | Confirmation that the temperature relaxation after stopping IR laser irradiation is due to heat dissipation

In this work, we assumed that IR laser irradiation creates a heat source within cells. We confirmed no changes that differed from this assumption when examining relaxation using this approach were detected. We here verified that our conclusions were affected by the possibility that (**a**) FPT responds to factors other than temperature during IR laser irradiation and/or (**b**) IR laser irradiation induces changes in molecular structure, distribution, or reactions (including physiological functions) without involving heat.

#### (a) Temperature selectivity of the FPT response during IR laser irradiation

During heating, factors other than temperature, such as viscosity, also change. Previously, we confirmed that the fluorescence response of FPT is temperature-selective and remains unaffected by factors such as pH, ionic strength, biomolecule concentration, and viscosity[1]. In this study, we investigated the temperature selectivity of the FPT response during IR laser irradiation

First, because FPT does not directly absorb IR laser light (Supplementary Figure 3), its response is independent of changes due to electronic excitation. Furthermore, CP having the same chemical composition as FPT but lacking temperature responsiveness did not respond to IR laser irradiation (Figures 2c, 3b, and Supplementary Figure 5). These results rule out the possibility that the FPT response is influenced by IR laser-induced excitation of intracellular molecules other than FPT, thermophoresis, or factors other than temperature (e.g., viscosity). The temperature selectivity of the FPT response demonstrated in these control experiments is supported by the results of Supplementary Figure 4. When we examined in detail the fluorescence lifetime of FPT and its component changes[2] reflecting the chemical environment (e.g., hydration, hydrogen bonding, and hydrophobicity), these properties during IR laser irradiation were identical to those during heating of the medium, where the energy input can be regarded as solely thermal.

These multifaceted control experiments indicate that changes in factors other than intracellular temperature, caused by heating within the physiological range, do not affect the FPT response.

**(b) Confirmation that the source of energy creating slow temperature relaxation is heat from IR laser**

First, we recorded the intracellular temperature relaxation when only heat was transferred to the cells by irradiating the extracellular medium with an IR laser. The results showed that intracellular temperature relaxation was significantly slower than that within liposomes (Supplementary Figure 17a), similar to the findings for intracellular heating (Figure 3f). This demonstrated that the intracellular temperature changes that we observed were due to heat delivered by IR laser.

Furthermore, the emergence of other exothermic reactions (e.g., metabolic heat production) induced by IR laser irradiation might influence the interpretation of temperature relaxation. Therefore, we verified that intracellular temperature relaxation after stopping IR laser irradiation is affected by intrinsic cellular heat production, including physiological functions. The results showed that slow temperature relaxation occurred even in cells with inhibited metabolism, thereby depleting energy and halting intrinsic cellular heat production (Supplementary Figure 17b). The results indicate that the intracellular temperature relaxation observed is due to heat dissipation rather than a time-dependent decrease in intrinsic heat production.

These experimental verifications from various perspectives confirm that temperature relaxation after stopping IR laser irradiation is due to heat dissipation.

### Supplementary Note 5 | Estimation of the temperature relaxation time in living cell based on heat conduction

The temperature relaxation time observed in cells was compared to that calculated with the assumption of heat conduction. The temperature change due to heat conduction after the supply of heat is stopped follows the heat conduction equation:

$$\frac{\partial(r\Delta T)}{\partial t} = \kappa \frac{\partial^2(r\Delta T)}{\partial r^2}; \qquad \text{(S3)}$$

$$\Delta T|_{t=0} = \begin{cases} \Delta T_0; 0 < r < a \\ 0; a < r \end{cases}$$

where $\Delta T$ is temperature, $c$ is specific heat, $\rho$ is density and $\lambda$ is thermal conductivity. $\kappa=\lambda/\rho c$ is thermal diffusivity. When the temperature in the region of radius $a$ is initially increased by $T_0$ K relative to the surroundings, the temperature relaxation after the disappearance of the heat source is given by the following exact solution:

$$\Delta T = \frac{\Delta T_0}{2r\sqrt{\pi\kappa t}} \left\{ \int_0^a \lambda e^{-\frac{(r-\lambda)^2}{4\kappa t}} d\lambda - \int_0^a \lambda e^{-\frac{(r+\lambda)^2}{4\kappa t}} d\lambda \right\} \qquad \text{(S4)}$$

Transforming equation (S4), we obtain

$$4\pi r^2 \Delta T = 2\pi r^2 \Delta T_0 \mathrm{Erf}\left[\frac{r-a}{\sqrt{4\kappa t}}, \frac{r+a}{\sqrt{4\kappa t}}\right] - 4r\Delta T_0\sqrt{\pi\kappa t}\left\{ e^{-\frac{(r-a)^2}{4\kappa t}} - e^{-\frac{(r+a)^2}{4\kappa t}} \right\} \qquad \text{(S5)}$$

$$\text{Note) } \mathrm{Erf}\,[x, y] \equiv \frac{2}{\sqrt{\pi}} \int_x^y e^{-\eta^2} d\eta.$$

From equation (S5), the average temperature in region $i$ ($\Delta T_{av,i}$) is defined as

$$\Delta T_{av,i}(t) \equiv \frac{\int_0^{r_i} 4\pi r^2 T dr}{\frac{4}{3}\pi r_i^3} \quad \text{(S6)}$$

The normalized average temperature of region $i$ ($\Delta T_{normal,i}$) is:

$$\Delta T_{normal,i}(t) \equiv \Delta T_{av,i}(t)/\Delta T_{av,i}(0) \quad \text{(S7)}$$

Using this equation (S7) and the values of thermal conductivity ($\lambda$: 0.618 W $m^{-1}$ $K^{-1}$), specific heat capacity ($c$: 4.178 J $g^{-1}$ $K^{-1}$) and density ($\rho$: 1 g $mL^{-1}$) of water, and the radius of heat source (a: 0.65 μm), we numerically calculated the time course of temperature. As shown in Supplementary Figure 18, the relaxation time was on the order of μs−ms. Meanwhile, because various biomolecules (e.g., proteins, lipids, and glycerol) are present in cells, we examined how their thermal properties, particularly their low thermal conductivity, influence the time required for temperature relaxation. We numerically calculated the time course of temperature by using the values of thermal conductivity ($\lambda$: 0.1 W $m^{-1}$ $K^{-1}$)[3,4], specific heat capacity ($c$: 3.9 J $g^{-1}$ $K^{-1}$)[5] and density ($\rho$: 1.05 g $mL^{-1}$)[6] of cellular components. As shown in Supplementary Figure 18, these comparisons revealed that, although the low thermal conductivity of cellular components slows temperature relaxation, the relaxation due to heat conduction still occurs on the order of ms. It should be noted that the temporal resolution of this temperature tracking method (9 ms) appears insufficient for detecting subtle differences in temperature relaxation caused by variations in the thermal properties of intracellular components.

### Supplementary Note 6 | The nature of the temperature change caused only by heat conduction

To understand the principle of intracellular temperature change, we examined the following two properties of heat conduction.

#### (a) Independence of the relaxation time of the average temperature on heating duration

The relaxation of temperature distribution in systems following equation (S3), with constant thermal diffusivity without a heat source term, only depends on the distribution of initial temperature. Therefore, when the regions defined for calculating averaged temperature (S6,7) are the same, the relaxation time of the average temperature ($\Delta T_{av}$) is identical and is not dependent on the heating duration. Note that according to the definition (S6), averaged temperature is equivalent to the amount of heat in the given averaged region; once the heat amount of the region is determined, the duration required for heating is irrelevant.

#### (b) Dependence of the relaxation time of the average temperature on area size

Next, we investigated the temperature change based on the nature of heat transfer by diffusion. In an aqueous solution, heat is dissipated to the surroundings by diffusion. To investigate the temperature change in water caused by heat transfer, we used the heat conduction equation (S3) and performed a numerical simulation of the temperature relaxation after stopping heating. Comparing the average temperature relaxation in regions of different sizes, we found that the time required for relaxation increases with the size of the observed region (Figure 4d).

### Supplementary Note 7 | Effect of thermal resistance at intracellular multiphase interfaces on heat dissipation rate

Within cells, organelles, phase-separated condensates, and locally concentrated biomolecules form complex multiphase interfaces. The interfacial thermal resistance created by numerous interfaces of this kind may slow heat dissipation. We examined the influence of these interfaces on intracellular heat dissipation rates, using interfacial thermal resistance values for lipid/protein–water interfaces obtained from experimental measurements or numerical analysis as a reference.

We used thermal resistance values for interfaces created by biomacromolecules estimated from theoretical calculations and experiments in previous studies. The thermal resistance ($R$) of a single interface is approximately 1–5×$10^{-9}$ $m^2$ K $W^{-1}$ (proteins: 3×$10^{-9}$ $m^2$ K $W^{-1}$; lipids: 1–5×$10^{-9}$ $m^2$ K $W^{-1}$)[3,7,8].

We assumed a situation in which the highest thermal resistance ($R$ = 5×$10^{-9}$ $m^2$ K $W^{-1}$) and total thermal resistance ($R_{total}$) of 1,000 interface layers within a representative length of $L$ = 10 μm hindered heat conduction within the cell. Considering the additional effect of interfacial thermal resistance on the intrinsic thermal conductivity ($\lambda$ = 0.1 W $m^{-1}$ $K^{-1}$) within the cell, the effective thermal conductivity ($\lambda_{eff}$) is:

$$R_{total} = \frac{L}{\lambda} + R \times 1000 = \frac{10 \times 10^{-6}}{0.1} + 5 \times 10^{-9} \times 1000 = 1.05 \times 10^{-4}\ \mathrm{m^2\ K\ W^{-1}}$$

$$\lambda_{eff} = \frac{L}{R_{total}} = \frac{10 \times 10^{-6}}{1.05 \times 10^{-4}} \fallingdotseq 0.095\ \mathrm{W}\ m^{-1}\ \mathrm{K^{-1}} \qquad \text{(S8)}$$

The presence of thermal resistance reduces the apparent thermal conductivity to 95% and extends the thermal relaxation time. However, the effect on thermal relaxation time is an increase of at most 5%, indicating that the order of magnitude remains unchanged (i.e., it significantly deviates from the measured temperature relaxation timescale in this experiment).

### Supplementary Note 8 | Estimation of intracellular temperature increase due to mitochondrial heat generation

Assuming that the dissipation of generated heat is entirely limited by the dominant energy conversion, we calculated the temperature increase due to spontaneous cellular heat generation using the measured intracellular heat dissipation time (approximately 1 s). Carbonyl cyanide 4-(trifluoromethoxy) phenylhydrazone (FCCP), an uncoupling agent in mitochondria, the most common target for measurement in intracellular thermometry, induces heat generation that lasts around 4.2 nW per cell[9]. When this heat remains in an adiabatic cell for 1 s, then the average temperature rise of whole cell can be estimated as follows:

$$\Delta T = \frac{Q}{c\rho V} = \frac{4.2 \times 10^{-9}}{3.9 \times 1.05 \times (0.9 \times 10^{-9})} = 1.1\ K \qquad \text{(S9)}$$

where $Q$ (= 4.2 nW × 1 s) is the amount of heat that remains in the cell, $c$ (= 3.9 J $g^{-1}$ $K^{-1}$) is the specific heat, $\rho$ (= 1.05 g $mL^{-1}$) is the density, and $V$ (= 30 μm × 30 μm × 1μm = 0.9 × $10^{-9}$ mL) is the volume of the cell. Note that this estimation assumes that this heat is shared by all of the cell volume based on the fact that the same dissipation rate and temperature change is recorded regardless of the chemistry of the thermometers, whereas the temperature change would be even larger if heat is shared by only some molecules.

### Supplementary Table

**Supplementary Table 1 | Coefficients in the polynomial of Equation (3)**

| | Living COS7 cell | COS7 cell extracts | Living HeLa cell |
|---|---|---|---|
| $A_1$ | 0 | 0 | 0.05049671042 |
| $A_2$ | 0.0603644271322865 | 0 | -2.266384469 |
| $A_3$ | -2.275549246 | 0.0434338143 | 42.0594063916615 |
| $A_4$ | 34.1786498914580 | -1.1433158570 | -412.714691229774 |
| $A_5$ | -255.3666680323850 | 10.9811316656 | 2256.40968069216 |
| $A_6$ | 949.582683193556 | -42.91691277 | -6509.582771 |
| $A_7$ | -1381.9478888330600 | 78.32179585 | 7755.40769741854 |

### Supplementary Figures

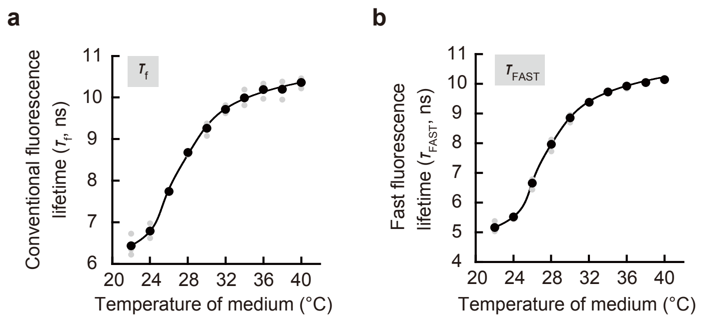


**Supplementary Figure 1 | Calibration curve of FPT for tracking of intracellular temperature changes.** The temperature-dependent conventional fluorescence lifetime ($\tau_f$, **a**) and fast fluorescence lifetime ($\tau_{FAST}$, **b**) of FPT in COS7 cells. Source data are provided as a Source Data file.

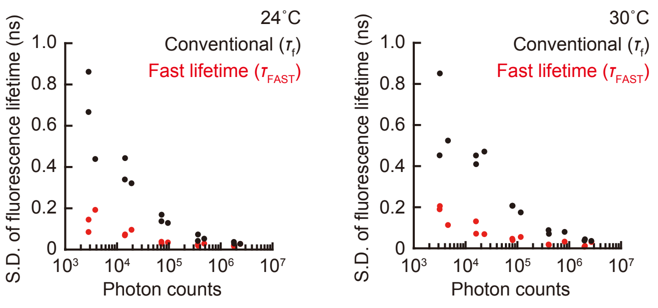


**Supplementary Figure 2 | Comparison of photon count-dependent fluorescence lifetime ($\tau_f$ vs. $\tau_{FAST}$) variation.** The fluorescence lifetime images of FPT were repeatedly obtained ten times and the SD of fluorescence lifetime ($\tau_f$ and $\tau_{FAST}$) of a single living COS7 cell was calculated with various accumulating durations corresponding to acquired photon counts. Two kinds of fluorescence lifetime ($\tau_f$ and $\tau_{FAST}$) at 24 °C and 30 °C are shown. Source data are provided as a Source Data file.

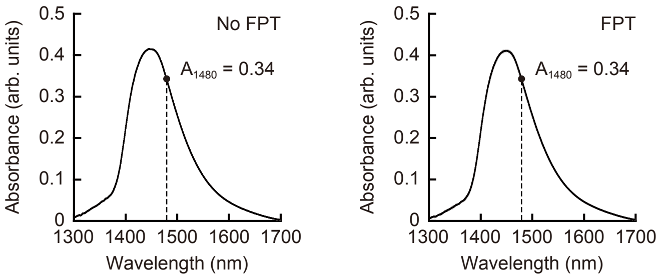


**Supplementary Figure 3 | Near-infrared absorption spectra of FPT solution.** The near-infrared absorption spectra of a KCl solution alone (150 mM) and that containing FPT (0.1 w/v%). The absorbance at 1480 nm ($A_{1480}$) is shown in the figures. Source data are provided as a Source Data file.

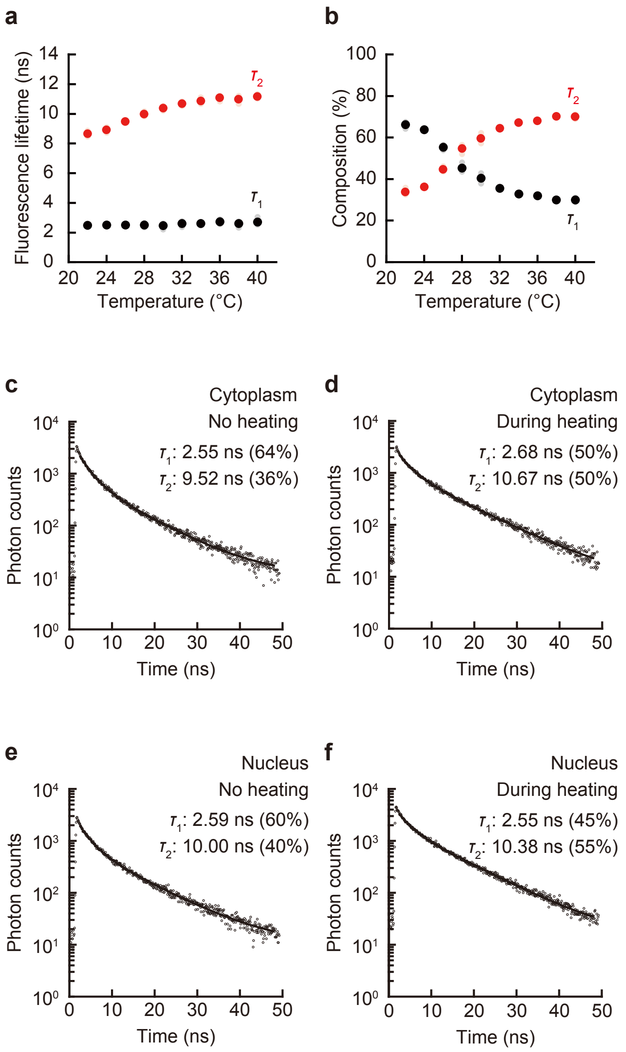


**Supplementary Figure 4 | Validation of the temperature increase in the area of IR laser heating based on the change in components (lifetime and composition) of fluorescence lifetime decay of FPT in living COS7 cells.** **a**,**b** The medium temperature-dependent two components of fluorescence lifetime decay of FPT in living COS7 cells. Note that these parameters are the values calculated in the analysis of conventional fluorescence lifetime. **a** Fluorescence lifetime–temperature diagram ($\tau_1$, shorter lifetime; $\tau_2$, longer lifetime). **b** Composition–temperature diagram of two components ($\tau_1$ and $\tau_2$). **c**–**f** Analysis of fluorescence lifetimes ($\tau_1$ and $\tau_2$) and their compositions of fluorescence decay curves of FPT without (**c**,**e**) and with IR laser heating (0.55 mW; **d**,**f**) in the cytoplasm (**c**,**d**) and in the nucleus (**e**,**f**). The changes in two components of fluorescence lifetime and composition were consistent with those of bulk temperature-dependent change in (**a**) and (**b**), confirming that there were no undesired chemical factors involved in the response of FPT. Source data are provided as a Source Data file.

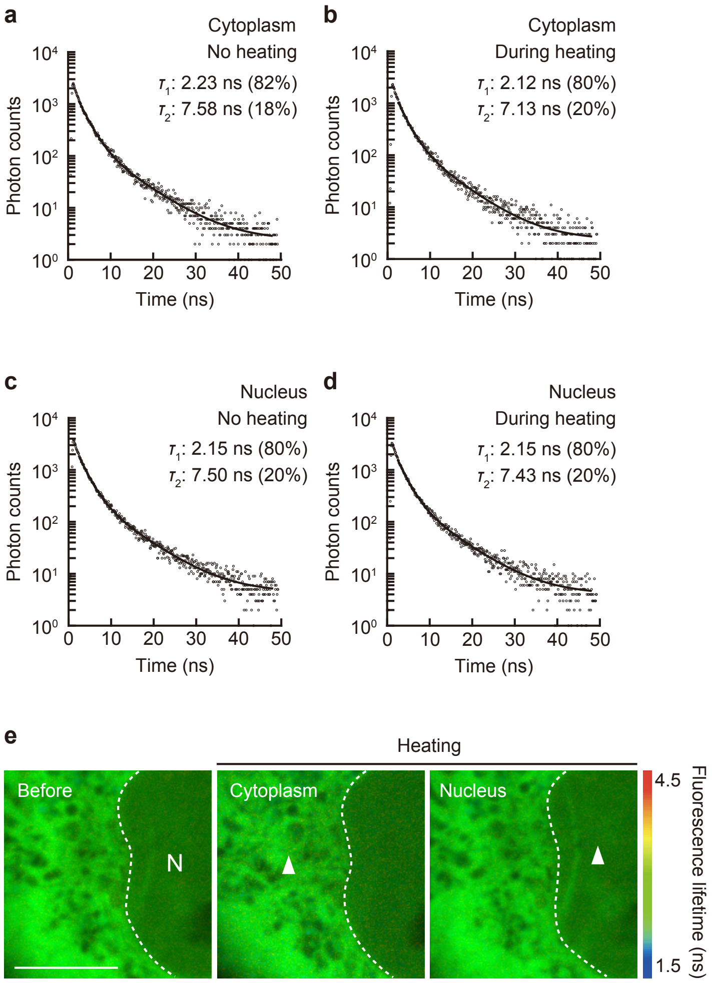


**Supplementary Figure 5 | Fluorescence decay curve and $\tau_{FAST}$ distribution of control copolymer (CP) in the IR laser irradiation region. a–d** Analysis of fluorescence lifetimes ($\tau_1$ and $\tau_2$) and their compositions of fluorescence decay curves of CP without (**a**,**c**) and with IR laser irradiation (0.55 mW; **b**,**d**) in the cytoplasm (**a**,**b**) and in the nucleus (**c**,**d**). **e** The $\tau_{FAST}$ distribution of CP during IR laser irradiation (0.55 mW) in the cytoplasm and in the nucleus. The arrowheads indicate the point of IR laser irradiation. Scale bar represents 10 µm. The experiment was performed three times with similar results. Source data are provided as a Source Data file.

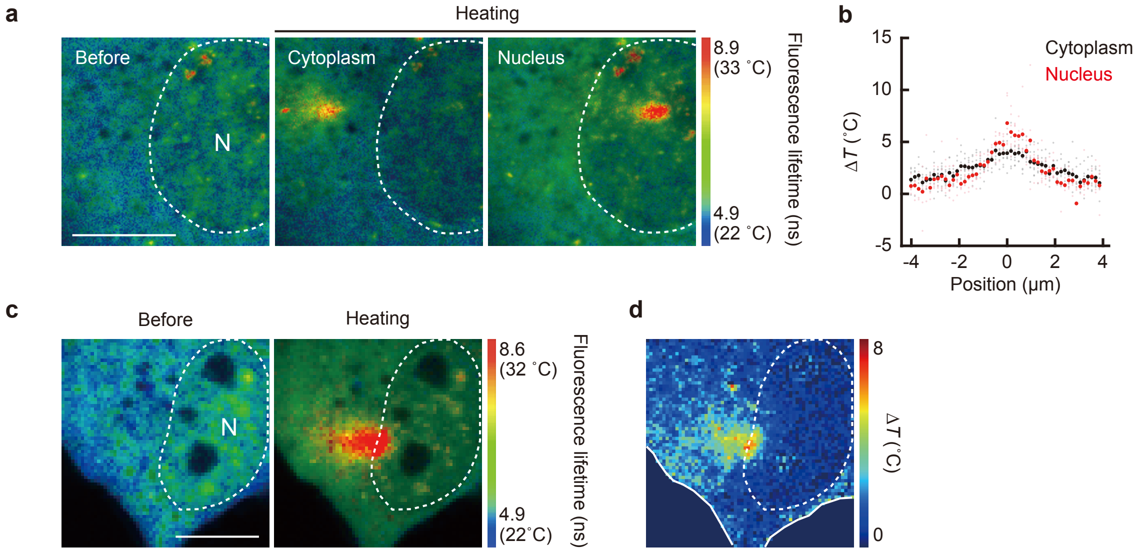


**Supplementary Figure 6 | IR laser heating of local regions in living COS7 cells**. **a** The steady-state temperature distribution during local heating (IR laser: 0.55 mW) in the cytoplasm and in the nucleus. **b** Line profiles of Δ*T* of the cytoplasm (*n* = 8 cells) and the nucleus (*n* = 6 cells). **c** The steady-state temperature distribution before and during local heating on the nuclear membrane. **d** The distribution of temperature increment upon heating (IR laser: 0.55 mW). Scale bars represent 10 μm. The experiments (**a**,**c**,**d**) were performed three times with similar results. Source data are provided as a Source Data file.

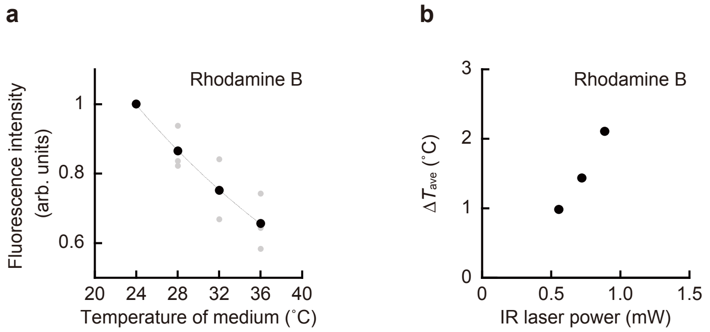


**Supplementary Figure 7 | Intracellular thermometry with Rhodamine B in living COS7 cells. a** The temperature-dependent fluorescence intensity decrease. An exponential approximation (Fluorescence intensity = $2.3153e^{-0.0351T}$, where *T* indicates the temperature; $R^2 = 0.99$) was used for the estimation of intracellular temperature change (Δ*T*). **b** The relationship between IR laser power and the temperature increase of whole cells (Δ*T*). Data are presented as mean ± s.e.m. (*n* = 10 cells). Source data are provided as a Source Data file.

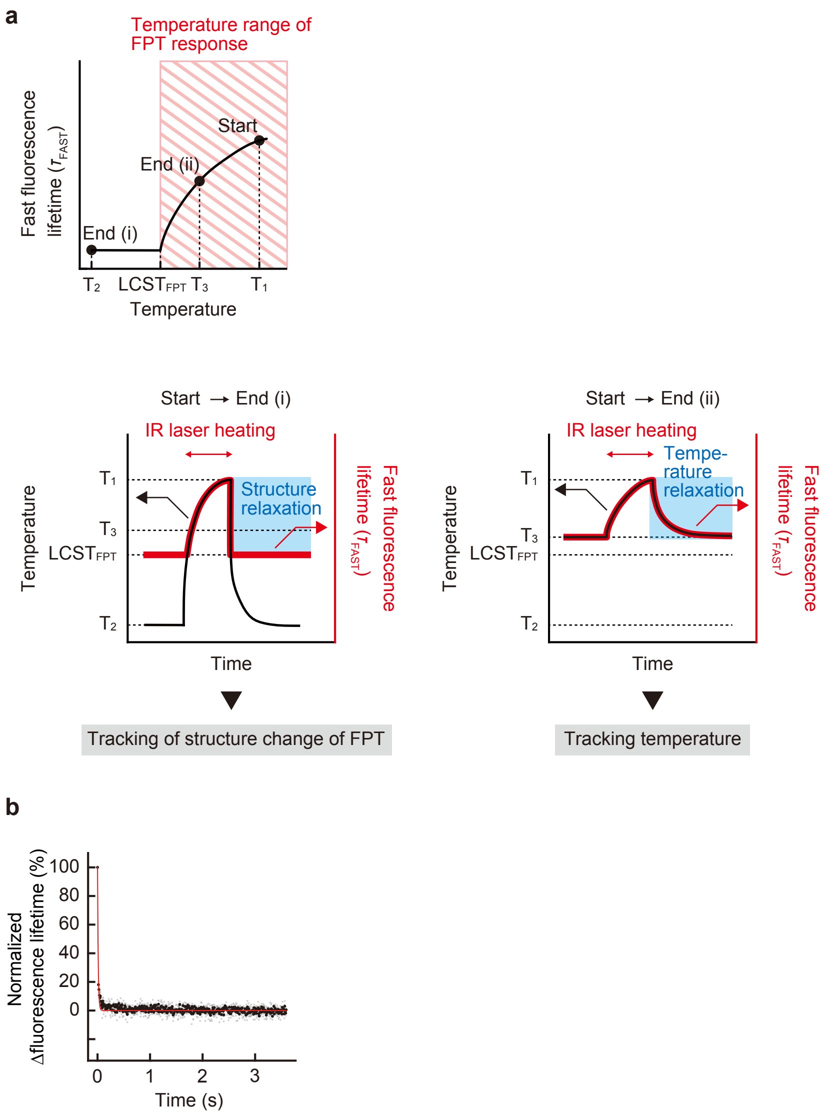


**Supplementary Figure 8 | The Examination of time for structural change of FPT in living COS7 cells. a** Schematic of this experiment. In principle, the temperature response dynamics of FPT includes both temperature and structural changes. FPT that changes its fluorescence lifetime due to the phase transition at temperatures above LCST does not respond at all below LCST (Upper panel). By lowering the base temperature before heating ($T_2$) below LCST, the response of FPT, which is sensitive only at temperatures above LCST, reflects structural relaxation (i, lower panel). On the other hand, when the base temperature before heating is set to $T_3$, which is higher than LCST, the fluorescence lifetime change reflects temperature (ii, lower panel). **b** The result for condition (i). The time required for FPT structural change was estimated by setting the temperature $T_2$ to a temperature (13 °C) that is significantly lower than the LCST of FPT (about 20 °C) to reduce the effect of temperature relaxation on the FPT fluorescence lifetime change. The red line shows an exponential approximation of the relaxation with a time constant of 13 ms. Source data are provided as a Source Data file.

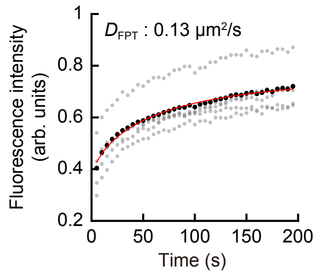


**Supplementary Figure 9 | Molecular diffusion of FPT in living COS7 cells.** Fluorescence recovery after photobleaching (FRAP) analysis of FPT. The recovery was fitted to the function by Soumpasis *et al.*[10] (red line) to yield a diffusion constant ($D_{FPT}$ = 0.13 µm$^2$ s$^{-1}$). Source data are provided as a Source Data file.

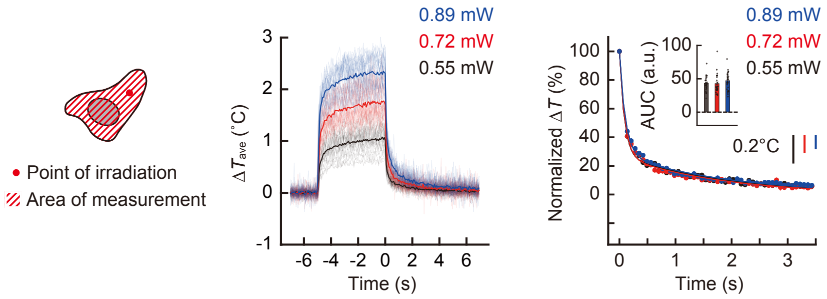


**Supplementary Figure 10 | Independence of temperature relaxation rate from temperature increase (Δ*T*).** Tracking of the average temperature change of whole cells when heated by continuous heating (5 s) with 0.55 (black), 0.72 (red), and 0.89 (blue) mW of IR laser irradiation (left). Normalized temperature relaxation of these three alternatives was compared (right). Scale bars represent indicated temperature changes. Data are presented as mean ± s.e.m. (*n* = 29 [0.55 mW], 30 [0.72 mW], and 30 [0.89 mW] cells). Inset in the right panel indicates area under the curve (AUC) of normalized temperature relaxation. The unit a.u. in AUC means arbitrary units. Source data are provided as a Source Data file.

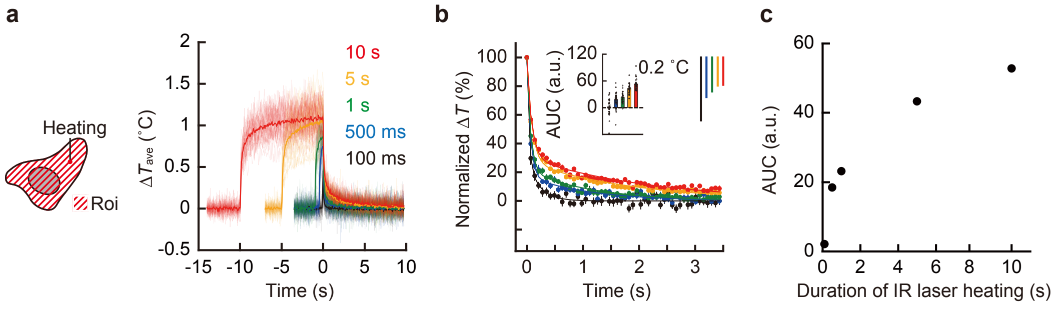


**Supplementary Figure 11 | The dependency of the intracellular temperature relaxation rate on the heating duration. a** Tracking of the average temperature change of whole cells when heated with an IR laser (0.55 mW) for 10 s (red), 5 s (yellow), 1 s (green), 500 ms (blue), and 100 ms (black). **b** Normalized temperature relaxation of these five alternatives was compared. Data including the inset are mean ± s.e.m. ($n$ = 30 [10 s], 29 [5 s], 30 [1 s], 30 [500 ms], and 28 [100 ms] cells). Inset indicates area under the curve (AUC) of normalized temperature relaxation. The unit a.u. in AUC means arbitrary units. Scale bars represent indicated temperature changes. **c** The relationship between the duration of heating and AUC of normalized temperature relaxation. Source data are provided as a Source Data file.

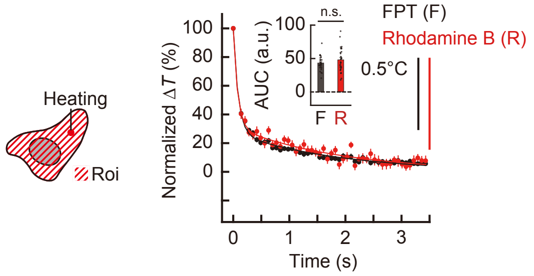


**Supplementary Figure 12 | Monitoring of temperature relaxation with Rhodamine B in living COS7 cells.** Comparison of normalized temperature relaxation after continuous heating with IR laser irradiation (5 s) between the result by FPT (black, Figure 3c) and that by Rhodamine B (IR laser: 0.55 mW, red). Scale bars represent indicated temperature changes. Data are presented as mean ± s.e.m. ($n$ = 30 [Rhodamine B] and 29 [FPT] cells). Inset in the right panel indicates area under the curve (AUC) of normalized temperature relaxation. The unit a.u. in AUC means arbitrary units. NS indicates not significant (Welch's two-sided $t$ test: $p = 0.216$). Source data are provided as a Source Data file.

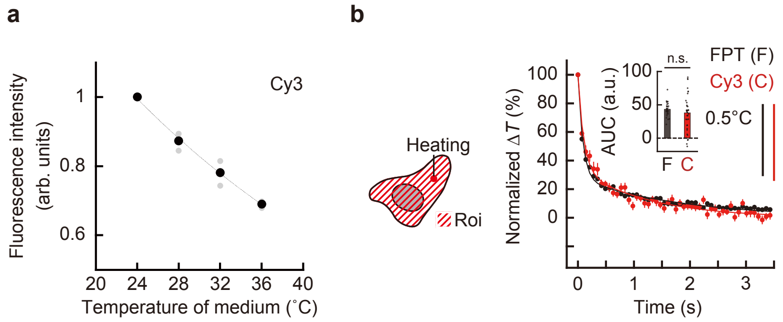


**Supplementary Figure 13 | Monitoring of temperature relaxation with Cy3 in living COS7 cells. a** The temperature-dependent fluorescence intensity decrease. An exponential approximation (Fluorescence intensity = $2.076e^{-0.0306T}$, where $T$ indicates the temperature; $R^2 = 0.99$) was used for the estimation of temperature change ($\Delta T$). **b** Comparison of normalized temperature relaxation after continuous heating with IR laser irradiation (5 s) between the result by FPT (black, Figure 3c) and that by Cy3 (IR laser: 0.55 mW, red). Scale bars represent indicated temperature changes. Data are presented as mean ± s.e.m. ($n$ = 35 [Cy3] and 29 [FPT] cells). Inset in the right panel indicates area under the curve (AUC) of normalized temperature relaxation. The unit a.u. in AUC means arbitrary units. NS indicates not significant (Welch's two-sided $t$ test: $p = 0.293$). Source data are provided as a Source Data file.

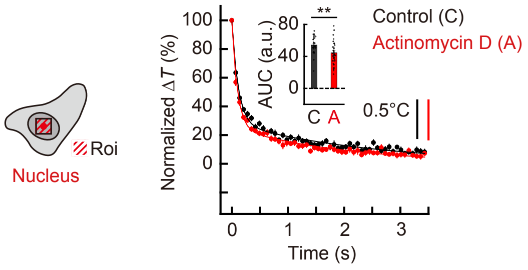


**Supplementary Figure 14 | The effect of transcription inhibition on the rate of temperature relaxation in the nucleus of living COS7 cells**. Normalized temperature relaxation of a square area with a 10 μm side centered on the heating point after continuous heating (IR laser: 0.55 mW for 5 s) in the nucleus was compared between untreated (control, black, Figure 3e) and Actinomycin D-treated (red) cells. Scale bars represent indicated temperature changes. Data are presented as mean ± s.e.m. ($n$ = 30 cells). Inset indicates area under the curve (AUC) of normalized temperature relaxation. The unit a.u. in AUC means arbitrary units. $^{**}P < 0.01$ (Welch's two-sided $t$ test: $p = 0.00486$). Source data are provided as a Source Data file.

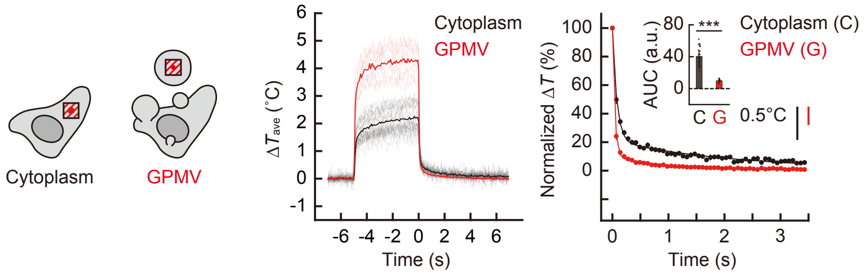


**Supplementary Figure 15 | The comparison of temperature relaxation rate between the cytoplasm of living COS7 cells and their vesicles containing a portion of the cytoplasmic components.** Tracking of the average temperature change of a square area with a 10 μm side centered on the heating point when heated with an IR laser (0.55 mW) for 5 s in the cytoplasm (control, black, Figure 3d) and in GPMV (red) of COS7 cells (left). Normalized temperature relaxation of these two alternatives was compared (right). Scale bars represent indicated temperature changes. Data are presented as mean ± s.e.m. ($n$ = 30 cells and 10 vesicles). Inset in the right panel indicates area under the curve (AUC) of normalized temperature relaxation. The unit a.u. in AUC means arbitrary units. $^{***}P <$ 0.001 (Welch's two-sided $t$ test: $p = 2.55\times10^{-17}$). Source data are provided as a Source Data file.

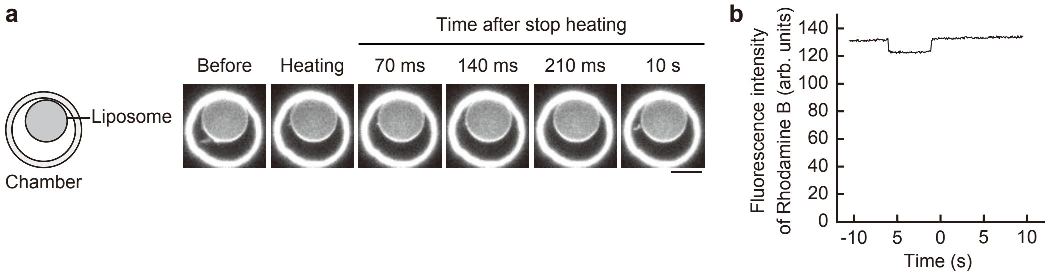


**Supplementary Figure 16 | Tracking of temperature in liposomes during transient heating. a** Fluorescence images of Rhodamine B in liposomes during heating (IR laser: 0.55 mW for 5 s). Liposomes were confined in a microchamber to prevent them from moving and disappearing from the field of view. The bright ring shape in the fluorescent images shows the structure of the chamber edge. Scale bar represents 10 μm. **b** Tracking temperature-dependent fluorescence intensity of Rhodamine B in liposomes. The experiment was performed three times with similar results. Source data are provided as a Source Data file.

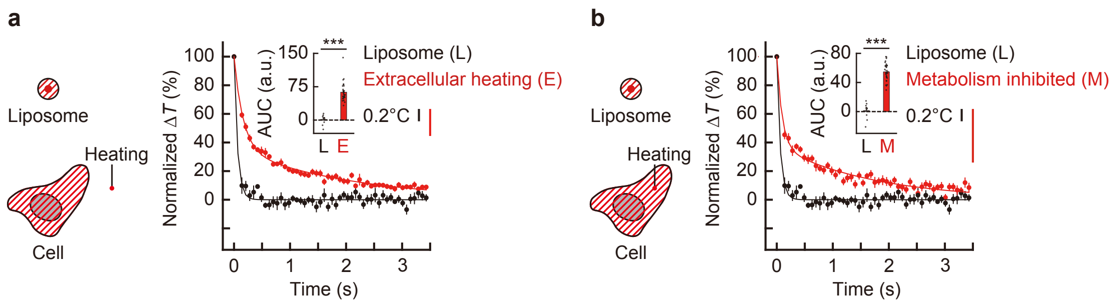


**Supplementary Figure 17 | The temperature relaxation rate in living cells when heating the extracellular space and when metabolic heat production is inhibited. a,b** Comparison of the normalized temperature relaxation in cells after cessation of heating of the extracellular medium (IR laser: 0.83 mW for 5 s, red; **a**) and of metabolic heat-inhibited cells (IR laser: 0.55 mW for 5 s, red; **b**), and the normalized temperature relaxation rate within liposomes (black, Figure 3f). Data are presented as mean ± s.e.m. ($n$ = 10 liposomes, 30 cells [extracellular heating], and 30 cells [metabolism inhibited]). Inset in the panels indicate area under the curve (AUC) of normalized temperature relaxation. The unit a.u. in AUC means arbitrary units. $^{***}P < 0.001$ (Welch's two-sided $t$ test: **a**, $p = 4.56\times10^{-13}$; **b**, $p = 2.42\times10^{-10}$). Source data are provided as a Source Data file.

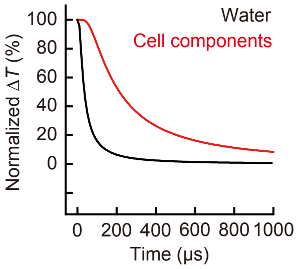


**Supplementary Figure 18 | Time-course of normalized temperature relaxation after stopping heating numerically simulated with the heat conduction equation.** The relaxations of average temperature of an area of a 5 μm radius centered on the heating point. The numerical simulations were performed with the heat conduction equation (S3) and the thermal properties reported for water (black) and cell components (red) (Supplementary Note 5). Source data are provided as a Source Data file.

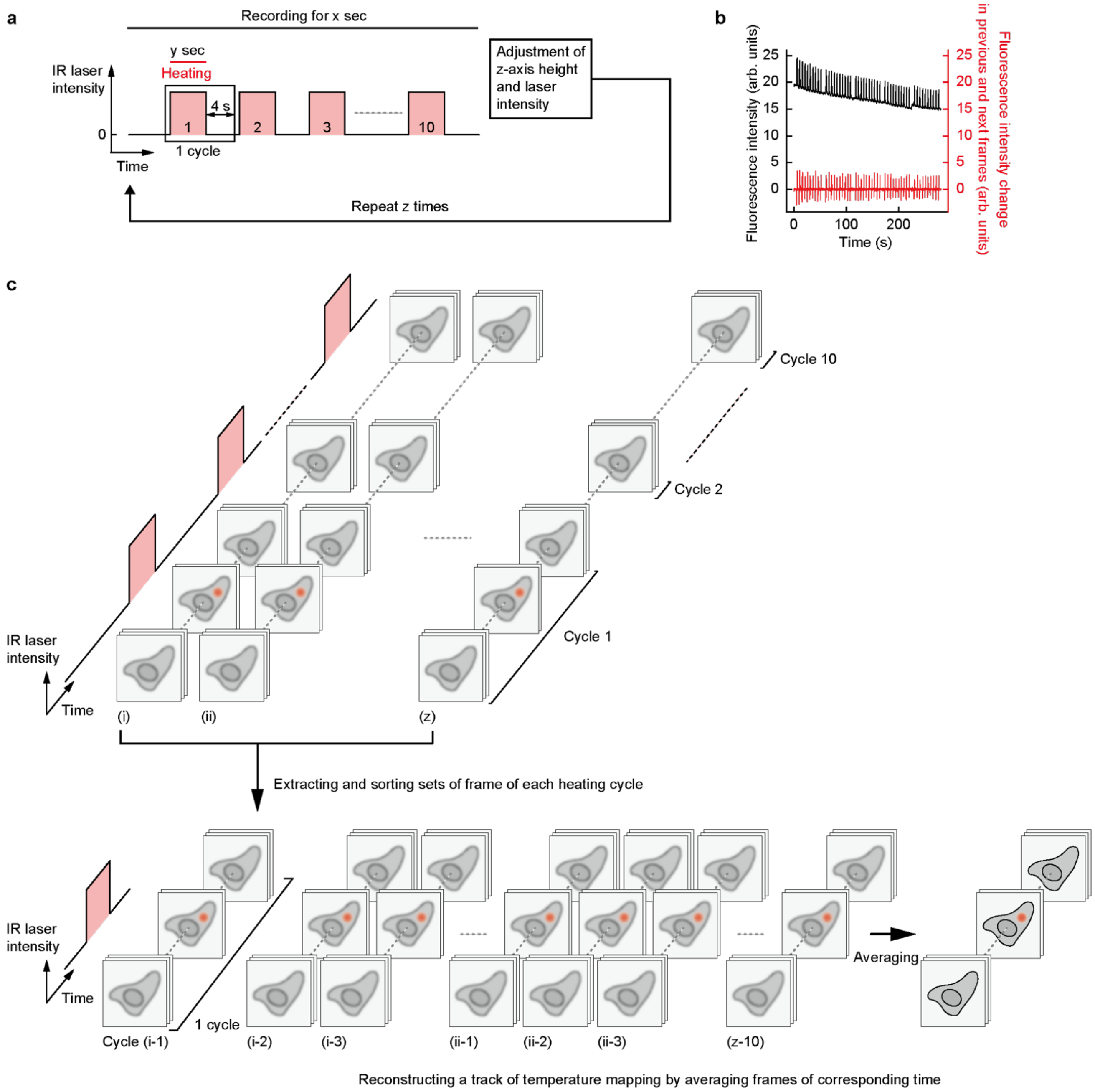


**Supplementary Figure 19 | Tracking of intracellular temperature distribution upon IR laser heating. a** Overview of imaging. Ten cycles of recording temperature relaxation for 4 s after y s of local heating in cells were repeated in continuous imaging for x s. **b** Time course of the fluorescence intensity in the field of view and the difference in fluorescence intensity between frames. **c** Video processing. The timing of heating stop was determined from the fluorescence intensity drop in b, and the subsequent time course of the fluorescence lifetime images was extracted for 4 s (one cycle). By averaging 10 × z movies for each cycle extracted from the continuous movie, a single high-resolution movie was reconstructed. Source data are provided as a Source Data file.

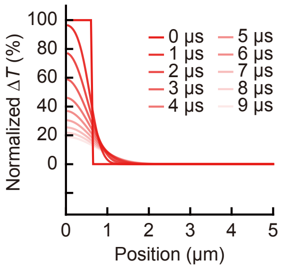


**Supplementary Figure 20 | Time-course of temperature distribution after stopping heating numerically simulated with the heat conduction equation.** Temperature distribution at each time after heat source term disappears. The numerical simulation was performed with the heat conduction equation (S3) and the thermal properties reported for cell components (Supplementary Note 5). Source data are provided as a Source Data file.

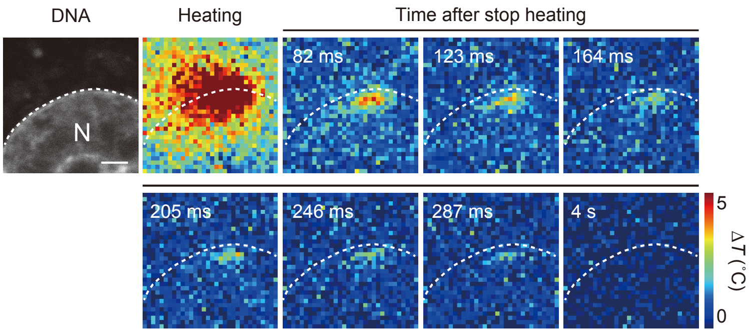


**Supplementary Figure 21 | Simultaneous mapping of nuclear and cytoplasmic temperature relaxation.** Real-time mapping of intracellular temperature immediately after stopping heating on the nuclear membrane (IR laser: 0.83 mW for 5 s). The nucleus was visualized with SiR-DNA. Scale bar represents 2 μm. Dotted line indicates the outline of the nucleus. The experiment was performed three times with similar results.

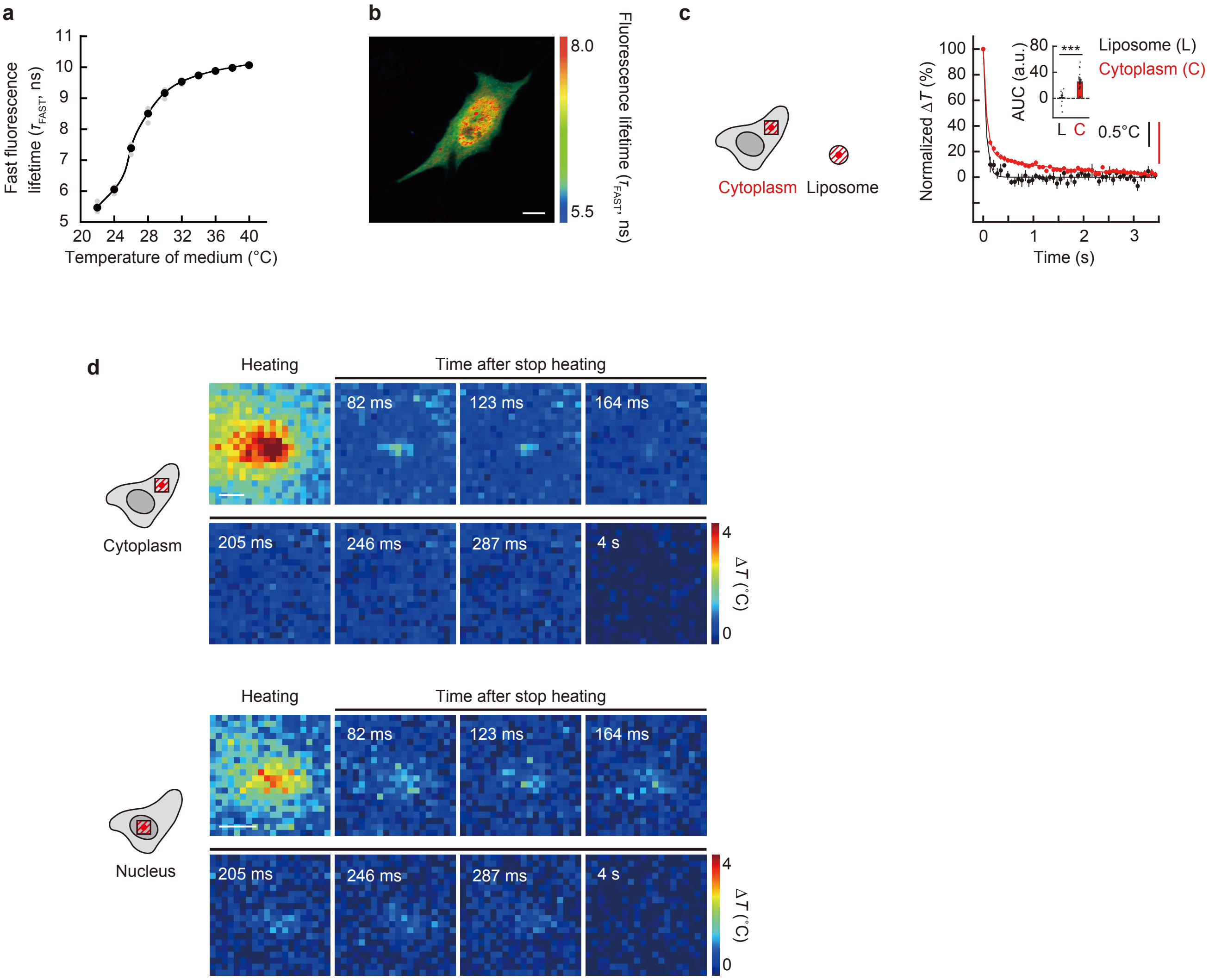


**Supplementary Figure 22 | Real-time mapping of temperature in living HeLa cell. a** The temperature response curve of FPT using fast fluorescence lifetime ($\tau_{FAST}$). **b** Fluorescence lifetime ($\tau_{FAST}$) images of FPT. Scale bar represents 10 µm. The experiment was performed two times with similar results. **c** Comparison of normalized temperature relaxation of liposome (Figure 3f, black) and that in a square area with a 10 µm side in living cells (red, IR laser: 0.55 mW) after the cessation of heating. Data are presented as mean ± s.e.m. ($n$ = 10 liposomes and 28 cells). Inset indicates area under the curve (AUC) of normalized temperature relaxation. The unit a.u. in AUC means arbitrary units. $^{***}P < 0.001$ (Welch's two-sided $t$ test: $p = 1.58\times10^{-5}$). **d** Real-time mapping of intracellular temperature in the cytoplasm (upper panels) and in the nucleus (lower panels) immediately after stopping heating (0.83 mW [cytoplasm] and 0.55 mW [nucleus] for 5 s). Scale bars represent 2 µm. The experiments were performed three (cytoplasm) and two (nucleus) times with similar results. Source data are provided as a Source Data file.

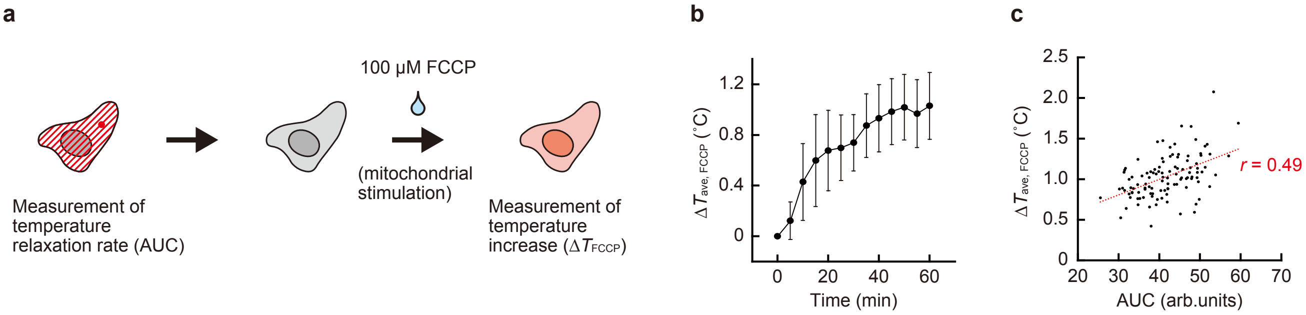


**Supplementary Figure 23 | The relationship between the rate of intracellular temperature relaxation and the temperature increase in living cells due to spontaneous heat generation. a** Experimental Schematic. The AUC of normalized temperature relaxation was quantified by IR laser heating (0.72 mW, 5 sec), followed by tracking the average temperature change of whole cells when mitochondria were stimulated with FCCP (100 µM) in the same cells. **b** Time course of the average temperature change in cells ($\Delta T_{FCCP}$) from immediately after the addition of FCCP. Data are presented as mean ± s.d. ($n$ = 109 cells). **c** The relationship between the AUC of normalized temperature relaxation and the temperature increase ($\Delta T_{FCCP}$) in the same cells. The average temperature increase of whole cells at 60 min after the addition of FCCP in 109 cells is plotted as a function of the AUC of normalized intracellular temperature relaxation. The least-squares regression line is shown by the red dotted line ($\Delta T_{FCCP}$ = 0.0192AUC + 0.2311, $r$ = 0.49). Source data are provided as a Source Data file.
